# Dopaminergic signalling is necessary, but not sufficient for cued fear memory destabilisation

**DOI:** 10.1101/564674

**Authors:** Charlotte R. Flavell, Jonathan L. C. Lee

## Abstract

Pharmacological targeting of memory reconsolidation is a promising therapeutic strategy for the treatment of fear memory-related disorders. However, the success of reconsolidation-based approaches depends upon the effective destabilisation of the fear memory by memory reactivation. Here, we show that the nootropic nefiracetam stimulates tone fear memory destabilisation to facilitate reconsolidation disruption by the glucocorticoid receptor antagonist mifepristone. Moreover, the enhancing effect of nefiracetam was dependent upon dopamine D1 receptor activation, although direct D1 receptor agonism was not sufficient to facilitate destabilisation. Finally, while the combined treatment with nefiracetam and mifepristone did not confer fear-reducing effects under conditions of extinction learning, there was some evidence that mifepristone reduces fear expression irrespective of memory reactivation parameters. Therefore, the use of combination pharmacological treatment to stimulate memory destabilisation and impair reconsolidation has potential therapeutic benefits, without risking a maladaptive increase of fear.

## Introduction

The disruption of memory reconsolidation represents a promising therapeutic approach for anxiety and trauma-related disorders. Pharmacological impairment of memory reconsolidation reduces fearful behaviour in rodents (Nader et al., 2000; Debiec et al., 2002), fear responses in experimental human studies (Kindt et al., 2009; Agren, 2014) and clinical symptoms in patients suffering with PTSD and phobias (Brunet et al., 2011; Soeter and Kindt, 2015).

While the efficacy of reconsolidation impairment appears relatively robust, targeting reconsolidation depends upon the success of destabilising the memory behaviourally via a memory reactivation session, which usually takes the form of cue re-exposure (Pineyro et al., 2013; Almeida-Correa and Amaral, 2014). It is increasingly evident that successful reconsolidation impairment is far from guaranteed (Kindt and van Emmerik, 2016), especially as there are unpredictable boundary conditions that govern memory destabilisation (Wideman et al., 2018). For example, we recently demonstrated that there appears to be no reliable basis upon which to predict the behavioural parameters that will trigger memory destabilisation/reconsolidation (Cassini et al., 2017). In fact, lack of replicability of reconsolidation impairments may well be due to poorly-understood boundary conditions on memory destabilisation (Bos et al., 2014).

This lack of reliability of memory destabilisation raises the potential that reconsolidation-disrupting pharmacological treatment might be applied to individuals with no chance of beneficial effect (because the memory has not been destabilised and so there is no reconsolidation process to impair). This has motivated the exploration of pharmacological enhancement of memory destabilisation (Bustos et al., 2010; Lee and Flavell, 2014; Gazarini et al., 2015; Ortiz et al., 2015). Here we further explored the potential to enhance the destabilisation of cued fear memories. In spite of recent promising results (Bustos et al., 2010; Lee and Flavell, 2014; Ortiz et al., 2015), we elected not to focus on D-cycloserine or ACEA as potentiators of destabilisation, partly due to the fact that D-cycloserine can enhance reconsolidation to strengthen fear (Lee et al., 2006) and there remains a degree of uncertainty concerning the effects of CB1 receptor modulation on fear memory reconsolidation (Lin et al., 2006; Ratano et al., 2014; Lee et al., 2019). Moreover, given the potential use of NMDA receptor and Cannabinoid CB1 receptor antagonists for the impairment of reconsolidation (Stern et al., 2012; Fattore et al., 2018), separable pharmacological targets for destabilisation enhancement and reconsolidation impairment would be desirable. Therefore, we focussed on additional mechanisms that have been implicated in memory destabilisation, starting with the demonstration that dopaminergic signalling in the amygdala is necessary for appetitive pavlovian memory destabilisation (Merlo et al., 2015). As a result, we tested whether dopamine D1 receptor agonism would enhance cued fear memory destabilisation. Moreover, we focussed on the use of the glucocorticoid antagonist mifepristone for the impairment of reconsolidation (Pitman et al., 2011), given our initial failure to replicate published findings with propranolol (Debiec and LeDoux, 2004).

## Methods

### Subjects

188 Lister Hooded rats (Charles River, UK; 200-225 g at the start of the experiment) were housed in quads under a 12 h light/dark cycle (lights on at 0700) at 21°C with food and water provided ad libitum apart from during the behavioural sessions. The cages were individually ventilated for the animals contributing to the data in Figs 1-3, and were standard cages for the animals contributing to the data in Fig 4 (due to a facility equipment change during the course of the project). The cages contained aspen chip bedding and environmental enrichment was available in the form of a Plexiglass tunnel. Experiments took place in a behavioural laboratory between 0830 and 1300. At the end of the experiment, animals were humanely killed via a rising concentration of CO2; death was confirmed by cervical dislocation. All procedures were approved by the University of Birmingham Animal Welfare and Ethical Review Body and conducted in accordance to the United Kingdom Animals (Scientific Procedures) Act 1986, Amendment Regulations 2012 (PPLs P8B15DC34 & P3B19D9B2).

**Fig. 1.**
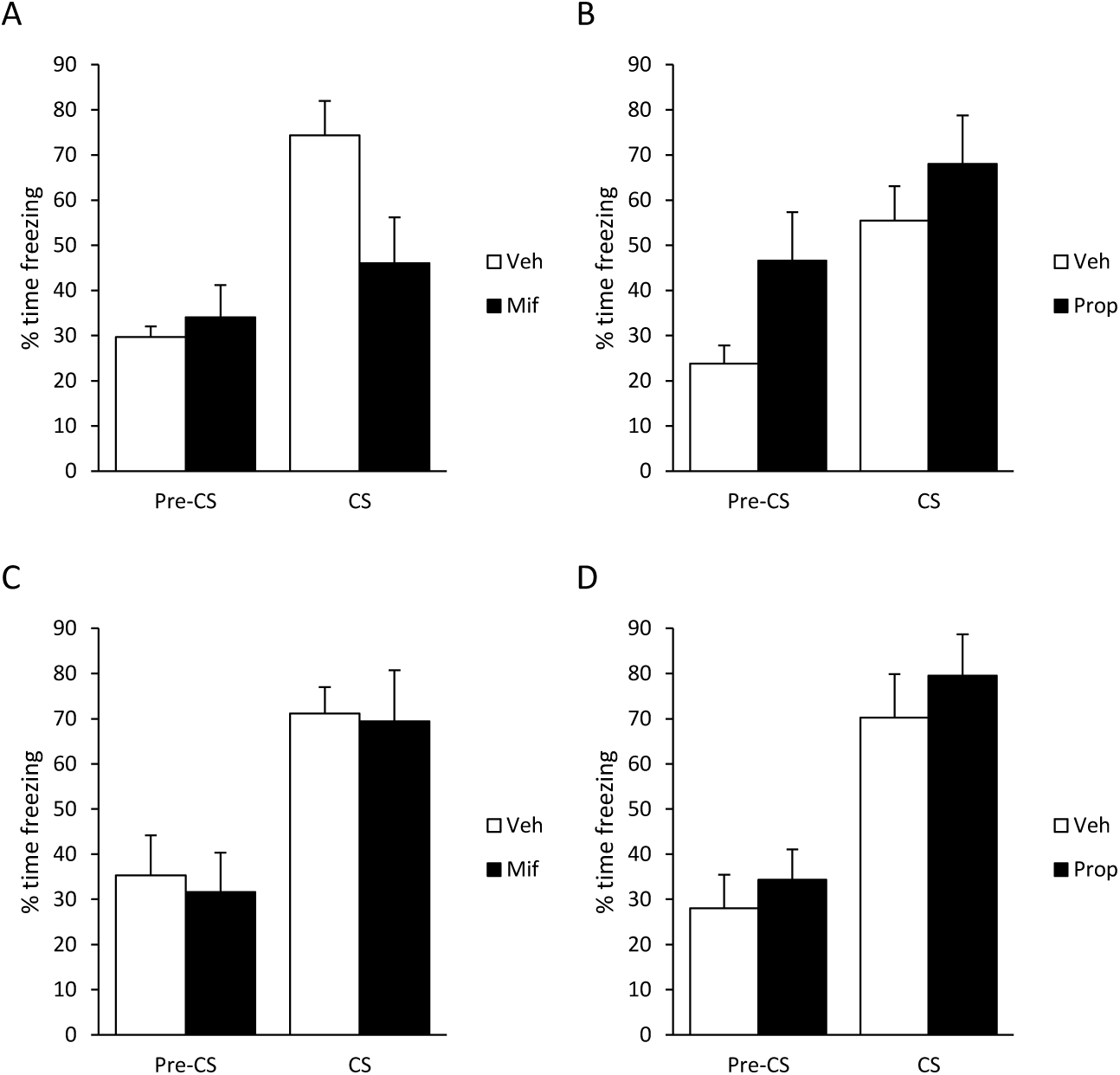
Systemic administration of mifepristone, but not propranolol impaired the reconsolidation of weak, but not strong, tone fear memory. After conditioning with a 0.5-s footshock, post-reactivation mifepristone (**A**), but not propranolol (**B**) impaired conditioned freezing to the tone, but not during the pre-CS period. After conditioning with a 1-s footshock, neither mifepristone (**C**) nor propranolol (**D**) impaired freezing during the pre-CS or tone periods. Data presented as mean + SEM.

### Drugs

All drugs were administered systemically at previously-established doses and timepoints. Mifepristone (Generon, UK) was injected at 30 mg/kg (60 mg/ml in propylene glycol, s.c.) immediately after memory reactivation (Pitman et al., 2011). DL-Propranolol (Sigma, UK) was injected at 10 mg/kg (10 mg/ml in saline, i.p.) immediately after reactivation (Debiec and LeDoux, 2004; Pitman et al., 2011). (+/-)-SKF38393 (Sigma, UK) was injected at 5 mg/kg (5 mg/ml in 5% DMSO in saline, i.p.) 5 min before reactivation (de Lima et al., 2011). Nefiracetam (Sigma, UK) was injected at 3 mg/kg (6 mg/ml in saline, i.p.) 1 hr before reactivation (Yoshii et al., 1997). SCH23390 (Tocris, UK) was injected at 0.1 mg/kg (0.1 mg/ml in saline, i.p.) 30 min before reactivation (Heath et al., 2015). Modafinil (Sigma, UK) was injected at 5 mg/kg (10 mg/ml in 50% DMSO in Saline, i.p.) 60 min prior to reactivation (Shanmugasundaram et al., 2015). Allocation to drug treatment was fully randomised within each experimental cohort of 8 rats.

### Behavioural equipment

The conditioning chambers (MedAssociates, VT) consisted of two identical illuminated boxes (25 cm × 32 cm × 25.5 cm), placed within sound-attenuating chambers. The box walls were constructed of steel, except by the ceiling and front wall, which were made of perspex. The grid floor consisted of 19 stainless steel rods (4.8 mm diameter; 1.6 mm centre-to-centre), connected to a shock generator and scrambler (MedAssociates, VT). Infrared video cameras were mounted on the ceiling of the chambers (Viewpoint Life Sciences, France) and used to record and quantify behaviour automatically.

### Behavioural procedures

Rats were conditioned and tested in pairs. They were initially habituated to the conditioning chamber for 1 hr. On the next day, they received a further 20-min habituation, followed by a single presentation of a single 30-s, 1.5-kHz tone, co-terminating with a 1-s (or 0.5-s), 0.4-mA footshock. There was a 2-min recovery period following the footshock delivery. 24 hours after training, the tone fear memory was reactivated by re-presenting the tone once for 60 s (the longer duration aiming to maximise prediction error, (Exton-McGuinness et al., 2015; Fernandez et al., 2016)), after a 60-s pre-CS period. 24 hrs after reactivation, conditioned freezing to the tone was assessed in a session identical to reactivation.

For the extinction experiment, all procedures were the same (with the 1-s footshock delivery) apart from the session 24 hours after training. Rats were exposed to ten 60-s tone presentations, after a 60-s pre-CS period and with 60-s intervals between each tone presentation (Lee et al., 2006).

### Statistical analyses

Data are presented as % time freezing (+ SEM) during the pre-CS period and tone presentation of the test. 9 subjects were excluded from the extinction experiment analyses due to equipment malfunction; 6 subjects were excluded as the primary endpoint was >2 s.d. from the group mean. The data were analysed in JASP (JASP Team, 2016) by repeated-measures ANOVA with Group and Phase (pre-CS vs. CS periods) as factors, followed by analyses of simple main effects of group at each phase. For the extinction experiment, the analysis used nefiracetam and mifepristone as separate factors in a 3-way repeated-measures ANOVA. Given the nature of the effects observed at test, additional analyses of the extinction session, as well as an exploratory ANCOVA (with performance at extinction included as the covariate) were conducted. The primary analyses were frequentist, with alpha=0.05 and either Cohen’s d or η^2^_p_ reported as an index of effect size, and the data were initially checked for normality. Significant group effects were explored with Tukey post-hoc pairwise comparisons. We also report BF_Inclusion_ and BF_10_ from parallel Bayesian analyses (Cauchy prior *r* = 0.707) as an estimate of posterior probability, with post-hoc tests as appropriate.

## Results

First, we showed that mifepristone, but not propranolol, was effective at impairing tone fear memory reconsolidation. Under weak single trial conditioning parameters (0.5-s, 0.4-mA footshock), immediate post-reactivation injection of mifepristone, but not propranolol impaired subsequent freezing to the conditioned tone at test (Fig. 1A, B). With mifepristone, there was a significant group x phase interaction (F(1,12)=16.0, p=0.002, η^2^_p_=0.57, BF_Inc_=28.0), with a simple main effect of group in freezing to the CS (t(12)=2.42, p=0.032, d=1.29, BF10=2.35), but not in the pre-CS period (t(12)=0.63, p=0.54, d=0.34, BF10=0.51). In contrast, with propranolol there no group x phase interaction (F(1,12)=0.66, p=0.43, η^2^_p_=0.05, BF_Inc_=1.74) and no main effect of group (F(1,12)=3.46, p=0.08, η^2^_p_=0.22, BF_Inc_=1.46). Moreover, planned analyses of simple main effects of group revealed no differences in freezing to the CS (t(12)=1.03, p=0.33, d=0.55, BF10=0.63). The disruptive effect of mifepristone was not replicated with stronger conditioning (1.0-s, 0.4-mA footshock, Fig. 1C, D). Post-reactivation injection of neither mifepristone nor propranolol had an effect on subsequent tone freezing. There were no group x phase interactions (mifepristone: F(1,12)=0.041, p=0.84, η^2^_p_=0.003, BF_Inc_=0.61; propranolol: F(1,12)=0.61, p=0.81, η^2^_p_=0.005, BF_Inc_=0.67) or main effects of group (mifepristone: F(1,12)=0.066, p=0.80, η^2^_p_=0.005, BF_Inc_=0.61; propranolol: F(1,12)=0.73, p=0.41, η^2^_p_=0.06, BF_Inc_=0.58). Planned analyses of simple main effects confirmed no group differences in freezing to the CS (mifepristone: t(12)=0.15, p=0.89, d=0.08, BF10=0.45; propranolol: t(12)=0.75, p=0.47, d=0.40, BF10=0.54). Therefore, our stronger conditioning parameters represent a boundary condition on tone fear memory reconsolidation, presumably under which our reactivation parameters were insufficient to destabilise the memory and render it vulnerable to the amnestic effect of mifepristone.

Next we tested whether pre-treatment with the D1R agonist SKF38393 would facilitate memory destabilisation and thereby render even the stronger tone fear memory vulnerable to disruption by mifepristone. The combination of pre-reactivation SKF38393 and post-reactivation mifepristone had no effect on test tone freezing compared to both SKF38393 + vehicle and vehicle + mifepristone (Fig 2A; group x phase: F(2,25)=0.13, p=0.88, η^2^_p_=0.01, BF_Inc_=0.19; group: F(2,25)=0.23, p=0.80, η^2^_p_=0.02, BF_Inc_=0.20; simple main effect of group on CS freezing: F(2,25)=0.03, p=0.97, η^2^=0.003, BF_Inc_=0.23). Numerical comparison with the previous groups receiving vehicle or mifepristone alone indicates that neither SKF38393 nor mifepristone in isolation had a disruptive effect on subsequent tone freezing.

**Fig. 2.**
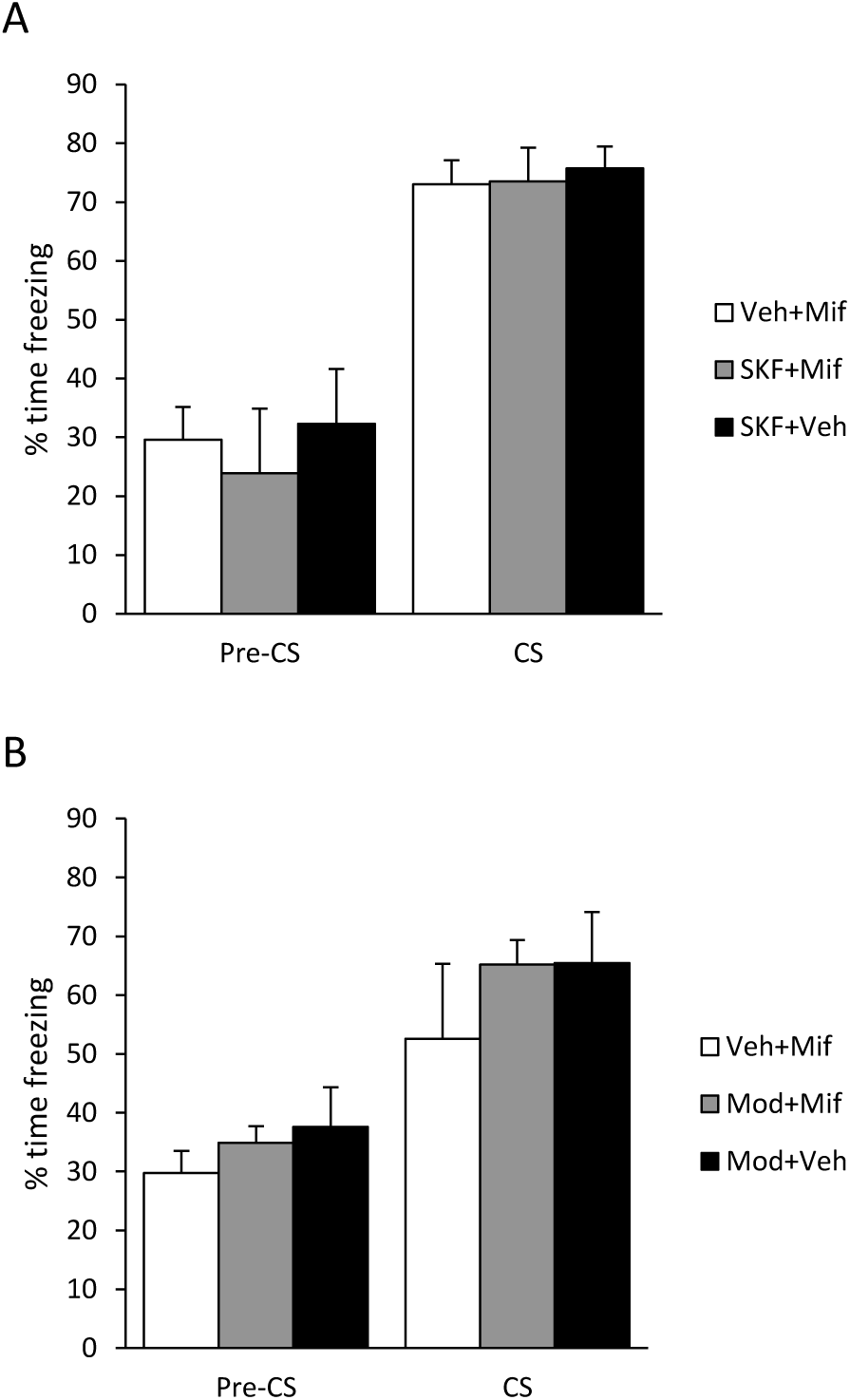
Enhancement of dopaminergic signalling did not stimulate fear memory destabilisation. After conditioning with a 1-s footshock, pre-reactivation SKF38393 (**A**) and modafinil (**B**) did not facilitate disruption of tone, or pre-CS, freezing by post-reactivation mifepristone, when compared to mifepristone and SKF38393 or modafinil alone. Data presented as mean + SEM.

We also tested whether the less-selective approach of pre-treatment with dopamine transporter blocker modafinil would facilitate memory destabilisation. The combination of pre-reactivation modafinil and post-reactivation mifepristone had no effect on test tone freezing compared to both modafinil + vehicle and vehicle + mifepristone (Fig. 2B; group x phase: F(2,19)=0.20, p=0.82, η^2^_p_=0.02, BF_Inc_=0.35; group: F(2,25)=1.06, p=0.37, η^2^_p_=0.10, BF_Inc_=0.37; simple main effect of group on CS freezing: F(2,25)=0.03, p=0.97, η^2^_p_=0.003, BF_Inc_=0.23). Numerical comparison with the previous groups receiving vehicle or mifepristone alone again indicated that neither modafinil nor mifepristone in isolation had a disruptive effect on subsequent tone freezing.

Given that selective agonism of D1 dopamine receptors or enhancement of dopaminergic neurotransmission did not appear to facilitate destabilisation, we adopted a broader spectrum approach, using the nootropic nefiracetam, which has effects on not only monoaminergic systems (Luthman et al., 1994), but also cholinergic signalling (Oyaizu and Narahashi, 1999) and calcium channels (Yoshii and Watabe, 1994), both of which have been implicated in memory destabilisation (Suzuki et al., 2008; Stiver et al., 2015). The combination of pre-reactivation nefiracetam and post-reactivation mifepristone reduced test freezing (Fig. 3A; group: F(2,18)=7.09, p=0.005, η^2^_p_=0.44, BF10=6.0; phase x group: F(2,18)=1.62, p=0.23, η^2^_p_=0.15, BF_Inc_=2.48). Analysis of simple main effects confirmed reduced freezing in the nefiracetam + mifepristone group to the tone compared to both nefiracetam + vehicle and vehicle + mifepristone (F(2,18)=, p=0.010, η^2^_p_=0.40, BF_Inc_=5.6; post-hoc p<0.05, BF10_(Nef+Mif vs Veh+Mif)_=2.2, BF10_(Nef+Mif vs Nef+Veh)_=4.1). The latter two groups froze at test at numerically higher levels to the vehicle and mifepristone groups in the previous experiment (nef+veh=88.5±3.3; veh+mif=85.2±5.8; veh=71.1±5.9; mif=69.4±11.3), suggesting again that neither nefiracetam nor mifepristone in isolation had a disruptive effect on subsequent tone freezing. Simple main effects revealed no significant effect of group on freezing in the pre-CS period (F(2,18)=3.01, p=0.074, η^2^_p_=0.25, BF10=1.4; post-hoc p=0.074, BF10_(Nef+Mif vs Veh+Mif)_=1.9, BF10(Nef+Mif vs Nef+Veh)=1.3). However, as there was no strong evidence for a selective effect on tone freezing, and poor evidence that nefiracetam + mifepristone did not impact upon pre-CS freezing, we conducted an exploratory ANCOVA, with pre-CS freezing as the covariate. This analysis confirmed the disruptive effect of nefiracetam + mifepristone on tone freezing (F(2,17)=4.23, p=0.032, η^2^_p_=0.33, BF_Inc_=4.5).

**Fig. 3.**
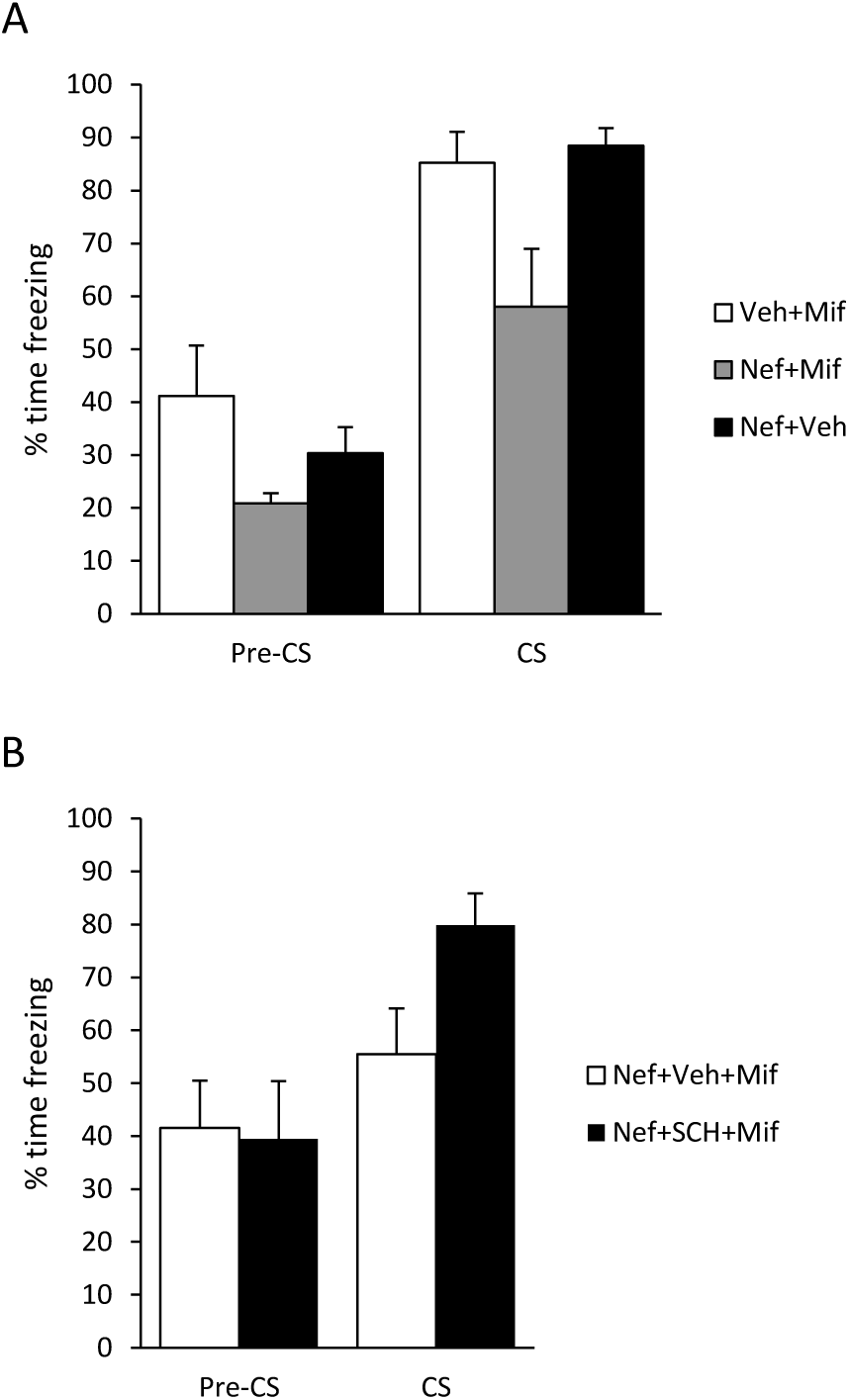
Systemic administration of nefiracetam stimulated fear memory destabilisation in a dopamine D1 receptor-dependent manner. After conditioning with a 1-s footshock, pre-reactivation nefiracetam facilitated disruption of tone, but not pre-CS, freezing by post-reactivation mifepristone, when compared to mifepristone and nefiracetam alone (**A**). When pre-reactivation treatment consisted of nefiracetam and SCH23390, mifepristone no longer impaired tone or pre-CS freezing (**B**). Data presented as mean + SEM.

While the mechanism of action of nefiracetam to facilitate memory destabilisation remains unclear, we focussed again on signalling at D1 dopamine receptors, testing whether such signalling is necessary for the enhancement of memory destabilisation. Co-pre-treatment with SCH23390 and nefiracetam blocked the facilitation of memory destabilisation. A nefiracetam-SCH23390-mifepristone group froze at higher levels at test relative to a nefiracetam-vehicle-mifepristone comparison group (Fig. 3B). There was a significant group x phase interaction (F(1,14)=5.96, p=0.029, η^2^_p_=0.30, BF_Inc_=4.1), with a simple main effect of group in freezing to the CS (t(14)=2.48, p=0.026, d=1.24, BF10=2.7), but not in the pre-CS period (t(14)=0.88, p=0.88, d=-0.08, BF10=0.43).

Given that the neurochemical mechanisms of destabilisation, reconsolidation and extinction overlap greatly, and that pharmacological approaches that impair reconsolidation can also disrupt extinction to maintain fear (Lee et al., 2006), we tested whether nefiracetam + mifepristone, or either drug individually, would affect extinction learning/consolidation. Nefiracetam and mifepristone were administered at the same timepoints relative to the extinction session as they had been in the previous reconsolidation experiments. There was an effect of mifepristone to reduce freezing to the tone, regardless of nefiracetam administration (Fig. 4A; phase x mifepristone: F(1,28)=7.92, p=0.009, η^2^_p_=0.22, BF_Inc_=10.9; phase x nefiracetam x mifepristone: F(1,14)=0.022, p=0.88, η^2^_p_=0.001, BF_Inc_=0.39). There was weak evidence that this effect of mifepristone on freezing to the tone was seen in both nefiracetam (phase x mifepristone: F(1,14)=4.36, p=0.056, η^2^_p_=0.24, BF_Inc_=1.54; mifepristone on tone freezing: t(14)=2.13, p=0.051, d=1.07, BF10=1.7; mifepristone on pre-CS freezing: (t(14)=0.10, p=0.92, d=0.05, BF10=0.43) and vehicle (phase x mifepristone: F(1,14)=3.71, p=0.075, η^2^_p_=0.21, BF_Inc_=1.88; mifepristone on tone freezing: t(14)=2.43, p=0.029, d=1.22, BF10=2.5; mifepristone on pre-CS freezing: (t(14)=0.086, p=0.93, d=0.04, BF10=0.43) conditions. Therefore, post-extinction mifepristone appears to reduce freezing to the tone CS.

**Fig. 4.**
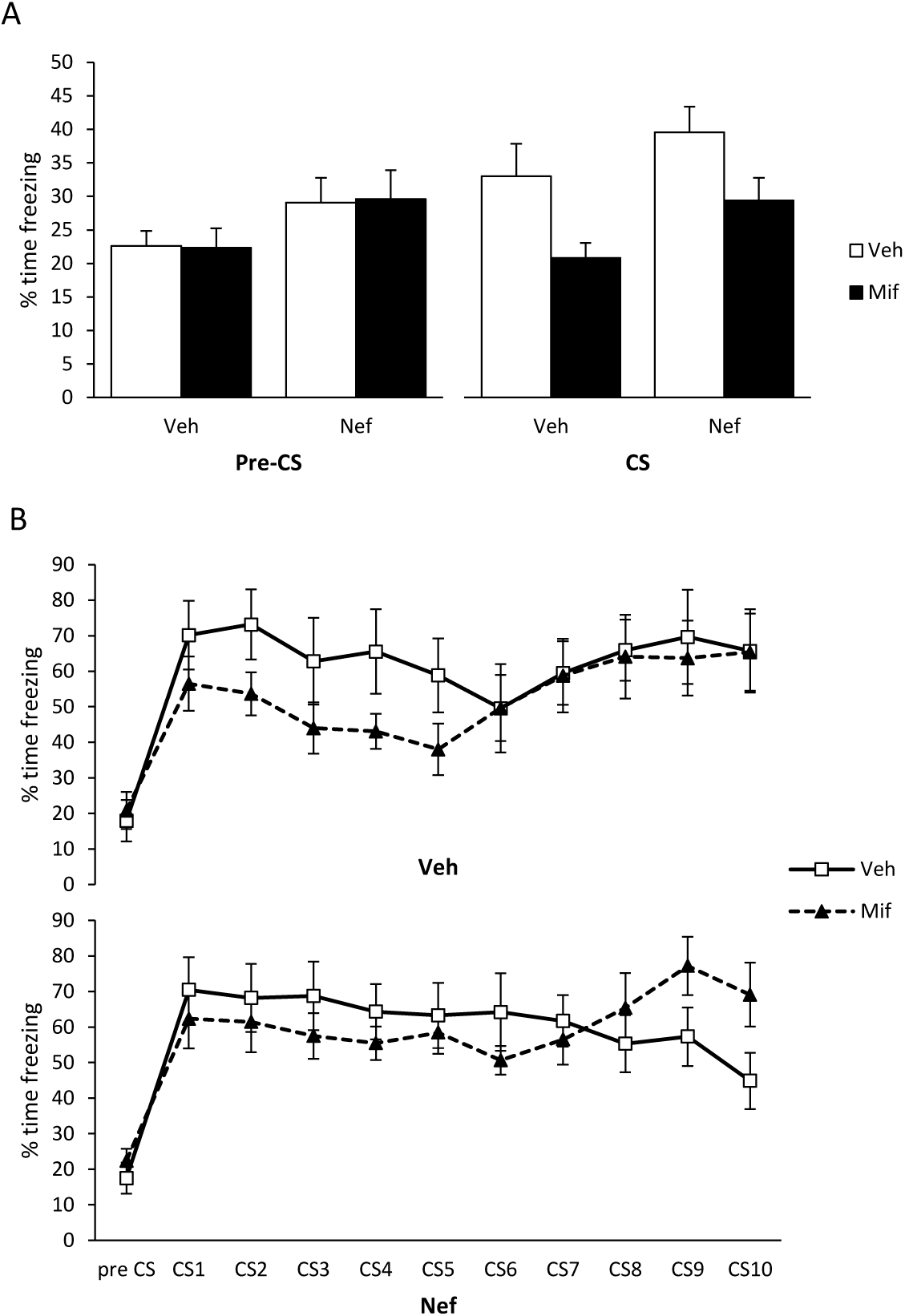
Effects of pre-extinction nefiracetam and post-extinction mifepristone. After conditioning with a 1-s footshock, nefiracetam was injected systemically prior to extinction and mifepristone immediately after extinction. At test, mifepristone reduced freezing to the tone, while nefiracetam increased freezing in both the pre-CS and tone periods (**A**). While there was no acute effect of pre-extinction nefiracetam at the extinction session, there was a pre-existing difference between the groups subsequently administered mifepristone compared to vehicle (**B**). Data presented as mean ± SEM.

We also observed an effect of nefiracetam to increase test freezing, irrespective of mifepristone administration (nefiracetam: F(1,28)=7.54, p=0.010, η^2^_p_=0.21, BF_Inc_=3.10; phase x nefiracetam: F(1,28)=0.031, p=0.86, η^2^_p_=0.001, BF_Inc_=0.69). This effect was observed across both tone (F(1,28)=4.80, p=0.037, η^2^_p_=0.15, BF_Inc_=1.85) and pre-CS (F(1,28)=4.75, p=0.038, η^2^_p_=0.15, BF_Inc_=4.03) periods. Further analysis suggested that the effect of nefiracetam on tone freezing was observed more clearly in mifepristone-(t(14)=2.27, p=0.039, d=1.14, BF10=2.05) than vehicle-treated rats (t(14)=1.14, p=0.28, d=0.57, BF10=0.65). Moreover, the effect of nefiracetam on pre-CS freezing was not obvious when the two subgroups were analysed independently (mifepristone: t(14)=1.50, p=0.16, d=0.75, BF10=0.89; vehicle: t(14)=1.61, p=0.13, d=0.80, BF10=0.98). Therefore, it remains unclear what is the major factor underpinning the elevation of freezing with nefiracetam.

Because the effect of nefiracetam appeared to occur whether or not mifepristone was subsequently administered, we checked whether pre-extinction nefiracetam had an acute effect at the extinction session that might have persisted to test (Fig. 4B). Analysis of the pre-CS period at the extinction session revealed no effect of nefiracetam or mifepristone (nefiracetam x mifepristone: F(1,26)=0.049, p=0.83, η^2^_p_=0.002, BF_Inc_=0.14; nefiracetam: F(1,26)=0.014, p=0.91, η^2^_p_=0.001, BF_Inc_=0.26; mifepristone: F(1,26)=0.77, p=0.39, η^2^=0.029, BF_Inc_=0.35). Analysis of freezing across the 10 tone presentations revealed no evidence for an acute effect of nefiracetam (tone x nefiracetam: F(2.6,66.6)=0.96, p=0.41, η^2^_p_=0.035, BF_Inc_=0.02; nefiracetam: F(1,26)=0.23, p=0.64, η^2^_p_=0.009, BF_Inc_=0.15). However, the analysis also revealed that there potentially were pre-existing differences at the extinction session between the groups subsequently administered with mifepristone (tone x mifepristone: F(2.6,66.6)=2.30, p=0.095, η^2^_p_=0.081, BF_Inc_=0.48; mifepristone: F(1,26)=0.89, p=0.36, η^2^_p_=0.033, BF_Inc_=0.30). Given that there appeared to be a small, albeit statistically non-significant, difference at the extinction session, we conducted an exploratory ANCOVA in order to determine whether the effect of mifepristone at test might be, at least in part, caused by pre-existing group differences. This analysis confirmed that, including freezing to the first tone at extinction as a covariate, there remained a significant effect of mifepristone (F(1,25)=7.06, p=0.014, η^2^_p_=0.22, BF_Inc_=5.14), as well as weaker evidence for an effect of nefiracetam (F(1,25)=4.62, p=0.041, η^2^=0.16, BF_Inc_=1.85).

Because the question of importance is whether the putatively-beneficial therapeutic administration of nefiracetam and mifepristone on destabilisation and reconsolidation might have alternative, and perhaps negative, effects if behavioural parameters promote extinction, we directly compared the nefiracetam + mifepristone group against the vehicle + vehicle group. There was weak evidence for a reduction in tone freezing (phase x group: F(1,14)=3.73, p=0.074, η^2^_p_=0.21, BF_Inc_=1.06; tone freezing: t(14)=0.65, p=0.53, d=0.33, BF10=0.49; pre-CS freezing: (t(14)=1.55, p=0.14, d=0.77, BF10=1.7 BF10=0.92). For consistency, we again conducted an exploratory ANCOVA, which confirmed no difference between the groups (F(1,11)=0.069, p=0.80, η^2^_p_=0.006, BF_Inc_=0.48). Therefore, the potentially beneficial effect of mifepristone and the contrasting negative effect of nefiracetam appear to interact with co-administration of the two drugs to result in no overall impact on freezing at test.

## Discussion

Our results show evidence that the combination of pre-reactivation systemic injection of nefiracetam and post-reactivation systemic mifepristone reduced fear expression to a fear conditioned tone. This disruptive effect was not observed following administration of either drug alone, or when nefiracetam was replaced by either the D1 dopamine receptor agonist SKF38393 or the dopamine receptor blocker modafinil. However, co-administration of the D1 dopamine receptor antagonist SCH23390 with nefiracetam and mifepristone eliminated the disruption of fear memory expression. The disruptive effect of nefiracetam and mifepristone was not replicated when an extinction session was used instead of memory reactivation. These results indicate that a combination treatment approach of nefiracetam to enhance memory destabilisation and mifepristone to impair reconsolidation may be effective for a reconsolidation-based treatment of fear memory disorders, without the risk of potentially counterproductive effects on extinction.

Systemic administration of mifepristone appeared to impair the reconsolidation of cued fear memoires under various conditions. A single re-exposure to an auditory stimulus is commonly used in reconsolidation studies (Nader et al., 2000), and we have previously used our current protocol to demonstrate that systemic administration of the NMDA receptor antagonist MK-801 impaired cued fear memory reconsolidation (Lee et al., 2006). Moreover, mifepristone has previously been shown to impair the reconsolidation of cued fear memories (Jin et al., 2007; Pitman et al., 2011), as well as a number of different memory types (Taubenfeld et al., 2009; Nikzad et al., 2011; Achterberg et al., 2014), although it has yet to be successfully translated to a human clinical setting (Wood et al., 2015). While we did not include a formal non-reactivation control condition (Dudai, 2004), the fact that mifepristone only disrupted freezing to the conditioned tone under certain parametric conditions rules out non-specific interpretations of the amnestic effect (see also Cassini et al., 2017). This boundary condition of initial conditioning strength has been previously observed across a number of settings (Suzuki et al., 2004; Rodriguez-Ortiz et al., 2005; Morris et al., 2006; Reichelt and Lee, 2012; Lee and Flavell, 2014). Importantly, it is not that strongly-learned memories cannot be triggered to undergo reconsolidation, but that the parameters of memory destabilisation are changed in a manner that is not easily predictable.

The failure of propranolol to impair fear memory expression at test under either of the two parametric conditions used here is somewhat surprising, given the previous evidence that propranolol does impair fear memory reconsolidation (Debiec and LeDoux, 2004; Kindt et al., 2009; Taherian et al., 2014; Ortiz et al., 2015; Villain et al., 2016). However, there are reports of failures to replicate the disruptive effect of propranolol in fear memories (Muravieva and Alberini, 2010; Pitman et al., 2011; Bos et al., 2014; Thome et al., 2016; Schroyens et al., 2017), as well as evidence that, at least in human studies, post-reactivation propranolol is less effective than pre-reactivation administration in impairing fear memory reconsolidation (Thomas et al., 2017). Given that mifepristone and propranolol had differential effects under identical parametric conditions, it is unlikely that the failure of propranolol here to disrupt fear memory reconsolidation represents a boundary condition on memory destabilisation. Therefore, it is perhaps more likely that the post-reactivation timing, and systemic injection nature, of propranolol administration explains the lack of disruptive effect. Alternatively, observations from studies of reconsolidation in other memory settings suggest that performance deficits following propranolol administration might be attributable to an attenuation of emotional value, rather than true memory impairment (Cogan et al., 2019), and there is a novel suggestion that b-adrenergic receptor signalling is necessary for destabilisation as well as reconsolidation (Lim et al., 2018). While it remains unclear why there was no evidence for an impairing effect of propranolol, the advantageous effect of mifepristone (see also Pitman et al., 2011) provided the basis for further exploration.

Under the stronger fear conditioning parameters, pre-reactivation systemic injection of nefiracetam rendered the post-reactivation administration of mifepristone effective in disrupting fear memory reconsolidation. The use of pre-reactivation pharmacological adjunctive treatment to facilitate reconsolidation impairments by other treatment has previously been demonstrated for stronger contextual fear memories (Lee and Flavell, 2014) and cued fear memories under conditions of ethanol withdrawal (Ortiz et al., 2015) and prior stress (Bustos et al., 2010). The common interpretation is that the additional pharmacological treatment facilitates memory destabilisation, rather than having an additive amnestic effect. Indeed the use of the cannabinoid CB1 receptor agonist ACEA (Lee and Flavell, 2014) and the NMDA receptor partial agonist D-cycloserine (Bustos et al., 2010; Ortiz et al., 2015) in previous studies was predicated on prior evidence that CB1 and GluN2B receptors are necessary for memory destabilisation (Ben Mamou et al., 2006; Suzuki et al., 2008; Milton et al., 2013).

The mechanism of action by which nefiracetam putatively enhances fear memory destabilisation remains somewhat unclear. The aforementioned clear bidirectional effects of CB1 and NMDA (GluN2B) receptor modulation on memory destabilisation (Szapiro et al., 2000; Ben Mamou et al., 2006; Suzuki et al., 2008; Lee and Flavell, 2014; Ortiz et al., 2015) have not been replicated here, in that the necessity for dopamine D1 receptor activation for cued fear memory destabilisation (Merlo et al., 2015) was not complemented here by any evidence that D1 receptor activation with SKF38393 is sufficient to enhance destabilisation. This is in spite of further evidence in the current study that D1 receptors are necessary for destabilisation under our experimental conditions. Co-administration of SCH23390 blocked the facilitative effect of nefiracetam, rendering mifepristone ineffective at impairing reconsolidation. This further suggests that D1 receptor activation is a necessary, but not the sole functional mechanism of action of nefiracetam to enhance destabilisation. It does not appear to be the case that the insufficiency of D1 receptor activation simply reflects the additional necessity of D2 receptor activation (Merlo et al., 2015), as the elevation of synaptic dopamine levels by modafinil-induced blockade of the dopamine transporter was similarly ineffective. This raises the question of whether nefiracetam acts up-or down-stream of D1 receptor activation. Acute administration of nefiracetam does elevate monoamine (including dopamine) levels under certain conditions (Luthman et al., 1994). However, nefiratecam also appears to augment intracellular memory-related processes to facilitate memory consolidation (Doyle et al., 1996; Nishizaki et al., 1998), raising the possibility that nefiracetam might enhance subthreshold intracellular destabilisation processes under boundary conditions of reconsolidation. The effect of co-administration of SCH23390 would then suggest that the subthreshold intracellular destabilisation results from an insufficient activation of D1 receptors, but this again is inconsistent with the failure of SKF38393 to enhance destabilisation.

The lack of effect of both SKF38393 and modafinil suggests there are non-dopaminergic mechanisms of action of nefiracetam. One highly likely additional mechanism of action is via L-type voltage-gated calcium channels (LVGCCs). Blockade of LVGCCs with systemic injections of nimodipine has been shown to prevent contextual fear memory destabilisation (Suzuki et al., 2008; Flavell et al., 2011; De Oliveira Alvares et al., 2013), and nefiracetam has pharmacological effects to enhance LVGCC calcium currents (Yoshii and Watabe, 1994). Therefore, we would predict that co-administration of nimodipine would replicate the effect of SCH23390 to prevent the enhancement of destabilisation by nefiracetam. A further possibility is that nefiracetam acts though cholinergic receptors, via an elevation of extracellular acetylcholine (Sakurai et al., 1998). While cholinergic receptors have not to our knowledge been studied in relation to fear memory destabilisation, activation of muscarinic acetylcholine receptors is sufficient to enhance destabilisation of object recognition memories (Stiver et al., 2015). Moreover, it is possible that activation of nicotinic acetylcholine receptors also contributes to object memory destabilisation (Stiver et al., 2015), and so the identified action of nefiracetam to elevate acetylcholine-induced currents at nicotinic acetylcholine receptors (Oyaizu and Narahashi, 1999) may contribute to the destabilisation of cued fear memories. However, perhaps the most likely mechanism of action is via NMDA receptors, given the effect of nefiracetam to potentiate NMDA receptor currents via interaction with the glycine binding site (Moriguchi et al., 2003), allied with the evidence that activation of NMDA receptors can facilitate destabilisation (Bustos et al., 2010; Ortiz et al., 2015).

Given the complex mechanistic relationship between destabilisation, reconsolidation and extinction (Almeida-Correa and Amaral, 2014; Merlo et al., 2014; Cassini et al., 2017), any potential therapeutic strategy that targets one of these processes has the potential to result in “off-target” effects on another process, leading to the possibility of maintaining or even enhancing the problematic memory (Lee et al., 2006; Tronson et al., 2006). Therefore, if the combination of pre-reactivation nefiracetam and post-reactivation mifepristone is to have genuine therapeutic promise, it is important to rule out potential counterproductive effects. This is especially the case, due to the observation that the parametric determinants of destabilisation and extinction are still not well understood (Merlo et al., 2014; Cassini et al., 2017; Merlo et al., 2018). Our results suggest that dual treatment with nefiracetam and mifepristone does not disrupt or facilitate cued fear memory extinction. Importantly, this lack of effect was observed under conditions that are appropriate for engaging extinction (Lee et al., 2006), and not due to the parameters of extinction training falling into the “null” or “limbo” space between destabilisation and extinction (Merlo et al., 2014; Cassini et al., 2017). This assumption is supported by the apparent effects of mifepristone and nefiracetam individually. However, these individual effects of mifepristone and nefiracetam indicate the need for caution when considering any translational application of the combined treatment.

The effect of nefiracetam to increase fear expression at test, and the suggestion that this increase in fear occurs even under conditions of mifepristone administration, indicates that pre-extinction nefiracetam disrupts extinction learning and/or consolidation. This is a novel observation, as to our knowledge the effects of nefiracetam on extinction of any memory have not previously been assessed. Such a disruption of extinction contrasts with the apparent facilitation of destabilisation, and as such may be inconsistent with the idea of a common labilisation system (Almeida-Correa and Amaral, 2014). However, it remains possible that nefiracetam modulates destabilisation and extinction via distinct mechanisms of action. Regardless of the mechanism of action, the fear-enhancing effect of nefiracetam alone raises concern that the therapeutic strategy of using nefiracetam to facilitate destabilisation might result in counterproductive effects on extinction.

In contrast, the apparent effect of mifepristone to reduce fear under conditions of extinction training further supports its potential benefit. A treatment that reduces fear expression irrespective of reactivation parameters would render that treatment less dependent upon understanding the boundary conditions of reconsolidation and extinction. However, it should be noted that the beneficial impact of mifepristone on extinction was rather modest, and it remains unlikely that mifepristone would have any effect in the “null point” between reconsolidation and extinction. Moreover, while infusion of mifepristone directly into the infralimbic cortex similarly enhanced the extinction of cued fear (Dadkhah et al., 2018), these results contrast somewhat with previous observations showing that intra-amygdala infusions of mifepristone did not directly affect extinction of fear-potentiated startle (Yang et al., 2006) and systemic injections of mifepristone did not affect the extinction of contextual fear (Ninomiya et al., 2010).

Ultimately, there is a need to explore the effects of mifepristone on extinction further, as well as determining the precise mechanisms of action of nefiracteam to enhance destabilisation and impair extinction. However, the present results support the premise that a strategy of enhancing destabilisation and impairing reconsolidation via dual drug treatment has the potential for reducing fear expression without risking fear potentiation.

